# Transcriptome Analysis of Human Induced Excitatory Neurons Supports a Strong Effect of Clozapine on Cholesterol Biosynthesis

**DOI:** 10.1101/2020.08.05.238212

**Authors:** Debamitra Das, Xi Peng, Anh-Thu Lam, Joel S. Bader, Dimitrios Avramopoulos

## Abstract

Antipsychotics are known to modulate dopamine and other neurotransmitters which is often thought to be the mechanism underlying their therapeutic effects. Nevertheless, other less studied consequences of antipsychotics on neuronal function may contribute to their efficacy. Revealing the complete picture behind their action is of paramount importance for precision medicine and accurate drug selection. Progress in cell engineering allows the generation of induced pluripotent stem cells (iPSCs) and their differentiation to a variety of neuronal types, providing new tools to study antipsychotics. Here we use excitatory cortical neurons derived from iPSCs to explore their response to therapeutic levels of Clozapine as measured by their transcriptomic output, a proxy for neuronal homeostasis. To our surprise, but in agreement with the results of many investigators studying glial-like cells, Clozapine had a very strong effect on cholesterol metabolism. More than a quarter (12) of all annotated cholesterol genes (46) in the genome were significantly changed at FDR<0.1, all upregulated. This is a 35-fold enrichment with an adjusted p = 8 ×10^−11^. Notably no other functional category showed evidence of enrichment. Cholesterol is a major component of the neuronal membrane and myelin but it does not cross the blood brain barrier, it is produced locally mostly by glia but also by neurons. By singling out increased expression of cholesterol metabolism genes as the main response of cortical excitatory neurons to antipsychotics, our work supports the hypothesis that cholesterol metabolism may be a contributing mechanism to the beneficial effects of Clozapine and possibly other antipsychotics.

## Introduction

Schizophrenia is a chronic debilitating psychiatric disorder that affects approximately 1% of population and is associated with poor quality of life and decreased life expectancy (McGrath et al., 2008). It is characterized by cognitive, positive (delusions, hallucinations, disorganized speech and thought) and negative (anhedonia, apathy, social and emotional withdrawal) symptoms (Owen et al., 2016). Schizophrenia is often diagnosed in early adulthood and in 39% of men and 23% of women, the disorder manifests by age 19 (Kao and Liu, 2010). Currently the primary treatment of schizophrenia is the use of antipsychotic medications, often in combination with psychosocial interventions and social support. The two leading theories for the etiology of schizophrenia, the glutamate and dopamine hypotheses, were both initially based on indirect evidence from the study of antipsychotic drugs (Howes et al., 2015), but are also backed by recent genetic findings (Consortium, 2014). Unfortunately, no single antipsychotic drug is effective for all patients. About half of those with schizophrenia will respond favorably to 1^st^ generation antipsychotics therapies including Chlorpromazine and Haloperidol. However 10-30% will not respond to 1^st^ or 2^nd^ generation antipsychotics, resulting in the persistence of positive symptoms (Ackenheil and Weber, 2004; Patel et al., 2014). Clozapine, once almost completely banished from the psychiatric pharmacopoeia, started being used again when its superior therapeutic effects in patients with refractory schizophrenia were demonstrated (Breier et al., 1999; Marder and Van Putten, 1988). In patients with schizophrenia who respond poorly to other antipsychotic medications, Clozapine, consistently improved both positive and negative symptoms as well as anxiety, mood, and hostile behaviors (Citrome et al., 2001; Lieberman, 1996; Suppes et al., 1999). But despite its efficacy, most clinical guidelines now recommend Clozapine monotherapy only to individuals with schizophrenia who have not adequately responded to at least two other antipsychotics. This is in part due to the potential risks associated with using Clozapine compared to other antipsychotics including agranulocytosis and adverse metabolic effects, such as weight gain, dyslipidemia, and hyperglycemia (De Hert et al., 2011). These metabolic side effects remain stumbling blocks in the use of Clozapine as a mainstream drug in today’s schizophrenia treatment that aim to improve patients’ symptoms as well as quality of life.

Cholesterol is an essential component for neuronal physiology. The brain is the most cholesterol-rich organ in the human body and possesses its own independent cholesterol metabolism machinery (Zhang and Liu, 2015) separate from that in peripheral tissues due to stringent diffusive properties of the blood-brain-barrier. Alteration in brain cholesterol metabolism have been associated with disorders such as Alzheimer’s, Parkinson’s, Huntington’s and amyotrophic lateral sclerosis (Jin et al., 2019). Cholesterol has been shown to have profound effects on the biophysical properties of neuronal membranes (Yeagle, 1985), thereby affecting the function of membrane-resident signaling components including ion channels, transporters and receptors (Barrantes, 1993; Lijnen, 1997; Spector and Yorek, 1985; Yeagle, 1989). Recent reports provide evidence that cholesterol is an essential element of the exocytosis apparatus and plays a crucial role in the biogenesis and transport of synaptic vesicles, whose membrane contains more cholesterol than other intracellular organelles (Pfrieger, 2003; Schmitz and Orso, 2001; Yeagle, 1985). Although neurons produce sufficient amount of cholesterol to survive, generate axons and dendrites and form a few inefficient synapses, it is the glial cells and astrocytes that supplement the neurons with the required additional cholesterol needed for massive formation of synapses (Pfrieger, 2003), while it has been shown that developing neurons produce higher amounts of cholesterol per cell than the astrocytes (Genaro-Mattos et al., 2019). It has been proposed that the effects of antipsychotics consistently identified in glial cells and in retinal pigment epithelia may reflect a mechanism of their action (Camargo et al., 2009; Ferno et al., 2006; Polymeropoulos et al., 2009; Vik-Mo et al., 2009).

With the goal to explore the effects of Clozapine on neuronal cells and choosing glutamatergic neurons inspired by the glutamate hypothesis for schizophrenia, we explored the effects of Clozapine at therapeutic levels on human glutamatergic neurons derived from iPSCs. We used changes in the transcriptome as a proxy to changes in the cell’s homeostasis which may reveal Clozapine’s therapeutic properties. Our findings add to our understanding of the potential therapeutic mechanisms of clozapine and how it may improve the symptoms of Schizophrenia.

## Material and Methods

### Induced pluripotent stem cell (iPSC) culture and maintenance

Human iPSC lines from normal controls were procured from the NIMH repository at Rutgers University and the cell line with ID MH0180967 (male of European descent) was used in this study (internal ID=RU01). Cells were cultured in StemFlex media (Gibco) on 6-well tissue culture plates coated with Laminin (Biolamina). Cells were dissociated with StemPro Accutase (Gibco) into single cell suspension and seeded in required density for the experiment. The Rock inhibitor Y-27632 dihydrochloride (Tocris) was added on the first day of passage at a concentration of 10uM. Cultured cells were tested to ensure they lack mycoplasma contamination.

### NGN2 lentivirus transduction

Lentiviruses expressing Ngn2 and rTTA were procured from the University of Pennsylvania Core facility (Lot# LV242 and LV243), which have been previously successfully used for induced neural differentiation (Zhang et al., 2013) to convert iPSCs into functional neurons with very high yield in less than 2 weeks. 250,000 RU01 iPSC cells were plated in each well of a 6 well plate and grown in Stem Flex media supplemented with Rock inhibitor. Ngn2 lentiviral infection using Polybrene (Santa Cruz) was done 24 hours post seeding. Briefly cells were fed with 2ml of fresh media and 2ul of Polybrene (1ug/ml) stock was added per well. In order to attain a multiplicity of infection (MOI) of 1-10 different volumes of both the Ngn2 and rTTA virus was added per well. In 4 of the 6 wells, leaving one as a polybrene-only control following amounts of EACH virus was added: 3ul, 5ul and 10ul. The virus infected cells were expanded and frozen stocks made for future differentiation. We selected the 10ul transduced Ngn2-RU01 cells for optimal neural differentiation.

### Neuronal differentiation of NGN2 transduced cells

250,000 Ngn2 transduced RU01 cells were plated on laminin coated 6 well plates **(Day -2)**. Cells were fed with fresh Stem Flex media the next day **(Day -1)**. Ngn2 expression was induced by Doxycycline on **Day 0** using an induction media consisting of DMEM/F12 (Thermo Fisher), N2 (Thermo Fisher), D-Glucose (Thermo Fisher), 2-βME (Life technologies), Primocin (Invivogen), BDNF (10ng/ml, Peprotech), NT3 (10ng/ml, Peprotech), Laminin (200ng/ml, Millipore Sigma) and Doxycycline (2ug/ml, Sigma). A puromycin selection was done on these cells on **Day 1,** 24 hours post Doxycycline induction using the same induction media supplemented with puromycin (2.5ug/ml). Surviving cells were harvested on **Day 2** and plated on either matrigel coated 24 well plates at a concentration of 100,000 cells/well for ICC studies or 1×10^6^ cells/well of a 6 well plate for drug exposure studies in neural differentiation media consisting of Neurobasal media (Thermo Fisher), B27 (Thermo Fisher), Glutamax (Thermo Fisher), Penn/Strep (Thermo Fisher), D-Glucose (Thermo Fisher), BDNF (10ng/ml), NT3 (10ng/ml), Laminin (200ng/ml) and Doxycycline (2ug/ml). Cells were fed with neural differentiation media every other day till Day 12. Cells were treated with 2uM Cytosineβ-D-arabinofuranoside hydrochloride (Ara-C) on Day 4 to reduce proliferation and eliminate non neuronal cells in the culture. Doxycycline induction was initiated at Day 0 and continued till Day 12 after which it was discontinued and cells were fed every two days thereafter till Day 21 with Neural maturation media consisting of Neurobasal media A (Thermo Fisher), B27, Glutamax, Penn/Strep, Glucose Pyruvate mix (1:100, final conc of 5mM glucose and 10mM sodium pyruvate), BDNF (10ng/ml), NT3 (10ng/ml), Laminin (200ng/ml) and Doxycycline (2ug/ml). Neurons were mature enough to be harvested by Day 21.

### Clozapine exposure and RNA extraction

The antipsychotic drug Clozapine (300ng/ml) was added to the neural maturation media on Day 18 at a concentration of 300ng/ml to match therapeutic plasma levels. Neurons were harvested at Day21 using Accutase for RNA extraction or fixed with PFA for ICC analysis. Six clones were exposed to Clozapine (diluted in methanol-CH3OH) and six clones to equal concentration of the solvent (methanol) and used as controls. Neuronal pellets were resuspended in lysis buffer and RNA extraction done using the Zymo Quick RNA Miniprep kit. Total RNA was sent to Novogene Corporation Inc. (Sacramento, CA) for rRNA depletion, library generation and RNA-sequencing.

### RNA sequencing analysis

Six clozapine exposed samples and four CH3OH control samples were included in the analysis after removing two CH3OH control for low library concentration or high degradation. The ten libraries were sequenced in one batch and all passed Novogene’s post-sequencing quality control. We received on average 28.2 million reads per sample with a maximum of 38 and a minimum of 23.3 million. The 150bp paired-end reads were aligned to human reference genome GRCH38 by Hisat2 version 2.1.0 (Kim et al., 2019). Samtools 1.9 (Li et al., 2009) produced corresponding BAM files and Stringtie 2.0.3 (Pertea et al., 2015) was used to assemble and estimate the abundance of transcripts based on GRCh38 human gene annotations (Pertea et al., 2016). Bioconductor package tximport computed raw counts by reversing the coverage formula used Stringtie with the input of read length (Soneson et al., 2015). The output then was imported to another Bioconductor package DESeq2 for differential gene expression analysis (Love et al., 2014). For visualization, R package pheatmap was used to produce the heatmap with dendrogram. The principal component analysis was conducted by Python package scikit-learn 0.22.1 (https://scikit-learn.org/stable/).

### Immunofluorescence studies

Neuronal cells in 24-well plates were washed once in PBS and fixed in freshly prepared 4% paraformaldehyde in PBS (pH 7.4) for 15 min. Cells were then permeabilized with PBS plus 0.1% Triton X-100 for 15 min. Cells were blocked in 10% goat serum for 1hr and stained with primary antibody at 4 °C overnight. The next day, cells were washed three times with PBS +0.1% Tween20 (PBST) and incubated with the appropriate secondary antibodies for 1hr, washed again thrice with PBST and counterstained with DAPI (Roche cat#10236276001, diluted 1:1000. Primary antibodies used were for neuronal markers MAP2 (mouse, Sigma cat# M1406, diluted 1:250), Synapsin (Rabbit, Sigma cat# S193, diluted 1:250), and TUJ1 (Mouse, Biolegend cat#801202, diluted 1:300).

## Results

### Establishing an IPSC-derived neuronal model to test the effects of Clozapine

Twenty-one days after doxycycline induction (Figure 1A), NGN2/rTTA-transduced human iPSC cells showed morphological progression to neurons in Bright field images (Figure 1B) as reported (Camargo et al., 2009) and immunofluorescence staining with cortical neuronal markers MAP2, Synapsin, and TUJ1 confirmed the neuronal identity (Figure 1C) and high differentiation efficiency which we attribute to the addition of Ara-C on Day 4 of culture reducing proliferation and eliminating all but post-mitotic cells in the culture (see methods). A series of neuronal and glutamatergic marker genes confirmed excitatory neuronal identity (Table 1).

**Table 1.**
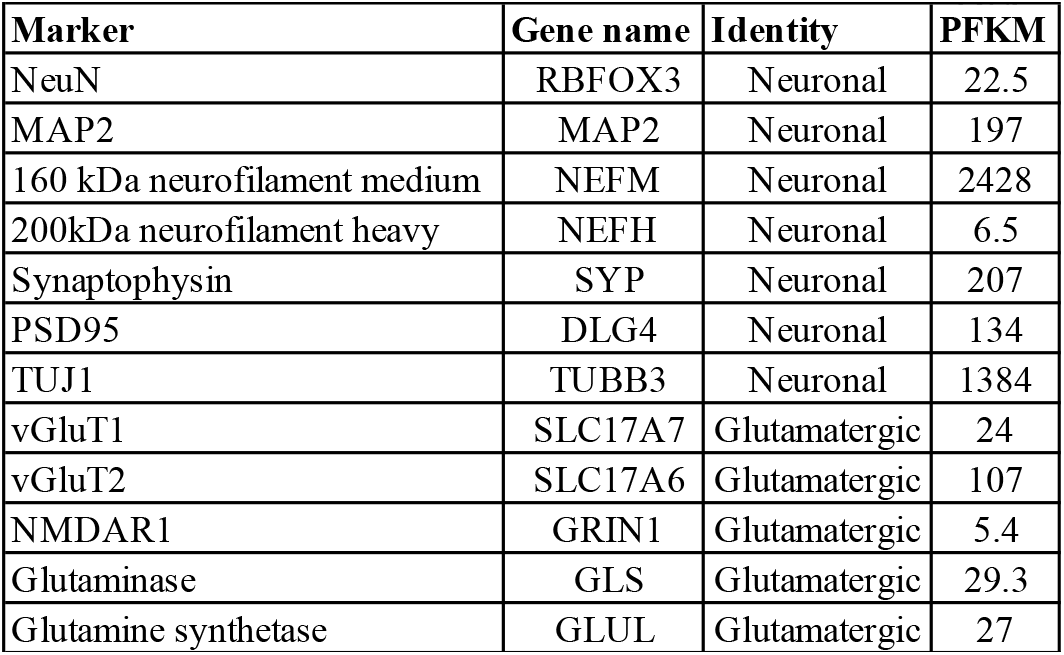
Differentiated iPSCs express excitatory neuronal markers

| Marker | Gene name | Identity | PFKM |
| --- | --- | --- | --- |
| NeuN | RBFOX3 | Neuronal | 22.5 |
| MAP2 | MAP2 | Neuronal | 197 |
| 160 kDa neurofilament medium | NEFM | Neuronal | 2428 |
| 200kDa neurofilament heavy | NEFH | Neuronal | 6.5 |
| Synaptophysin | SYP | Neuronal | 207 |
| PSD95 | DLG4 | Neuronal | 134 |
| TUJ1 | TUBB3 | Neuronal | 1384 |
| vGluT1 | SLC17A7 | Glutamatergic | 24 |
| vGluT2 | SLC17A6 | Glutamatergic | 107 |
| NMDAR1 | GRIN1 | Glutamatergic | 5.4 |
| Glutaminase | GLS | Glutamatergic | 29.3 |
| Glutamine synthetase | GLUL | Glutamatergic | 27 |

**Figure 1.**
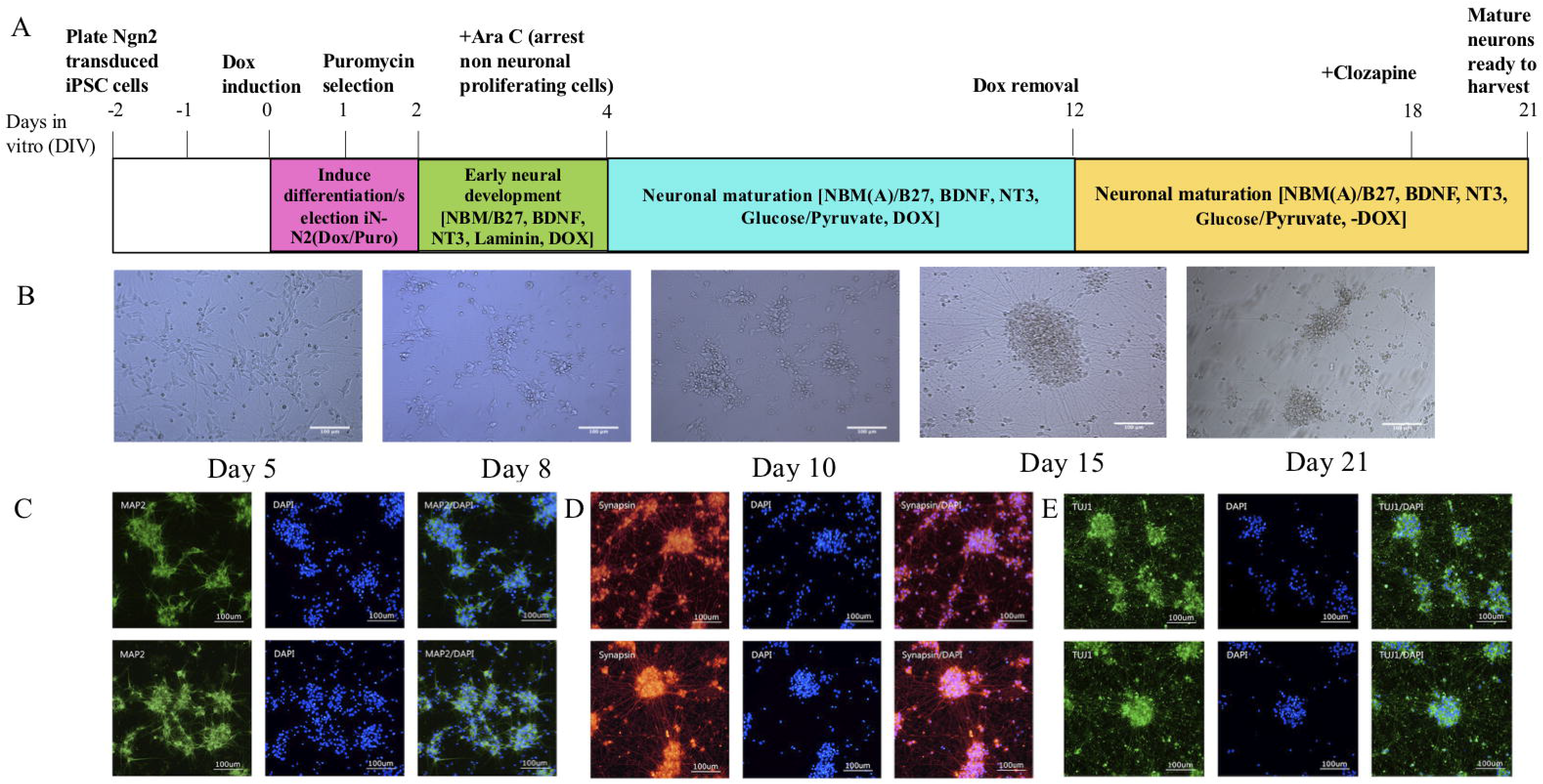
Experimental Timeline. (A) This figure outlines the timeline from plating Ngn2 transduced human iPSC cells through induction, differentiation, drug exposure and mature neuronal est. (B) Bright field images showing the morphological progression ofiPSC cells when induced to differentiate into cortical excitatory neurons using NGN2/rTIA (C-E) luorescence staining ofinduced neurons with cortical neuronal markers MAP2, Synapsin, and TUJI, respectively.

### Differential expression analysis between Clozapine treated and untreated neurons

Two of the control lines did not pass QC and were dropped from analysis. Genes with low read counts (< 10) were removed from analysis to decrease noise, leaving 19, 296 genes for analysis. Differential expression analysis between Clozapine exposed and unexposed neurons showed 51 genes changed at FDR <0.05, 122 genes at FDR <0.1 and 286 genes at FDR<0.2. The list of all genes and the analysis results are shown in Supplementary Table 1. We then used the Panther bioinformatics platform (www.pantherdb.org) to identify enrichment in functional categories among the differentially expressed genes at all 3 levels of significance. Functional annotation was available for 16,135 of the 19,296 genes. Highly significant enrichment for cholesterol biosynthesis genes was observed at all three significance level cutoffs reaching 35-fold at FDR<0.1 including 12 of the 46 annotated cholesterol biosynthesis genes in the entire gene set (enrichment FDR = 1.70E^−10^). All 12 cholesterol genes where up-regulated by the exposure to Clozapine (Figure 2C, 2D). Notably, no other functional category approached statistically significant enrichment. A heatmap of the set of all 44 cholesterol genes is shown in Figure 2B, the significantly changing genes marked with an asterisk. Exposed cells showed more variability in expression, likely reflecting pipetting errors and differences in cell confluence affecting the effective drug dose. Most differences were observed in the genes with higher expression and the corresponding dendrogram perfectly separated exposed from unexposed cells. Given this last observation, and to gage the magnitude of the effect of Clozapine on the entire set of cholesterol metabolism genes, we ran a principal component analysis using the entire set of 46 annotated cholesterol genes. This analysis showed that PC1, capturing the largest fraction of variance in cholesterol genes, perfectly separated exposed from unexposed cells (Figure 2A), underlining the profound effect of Clozapine on neuronal cell cholesterol metabolism. In fact, on Table 2 which lists the results for all cholesterol genes in the dataset it can be seen that most genes that did not achieve significance were also upregulated by Clozapine.

**TABLE 2:**
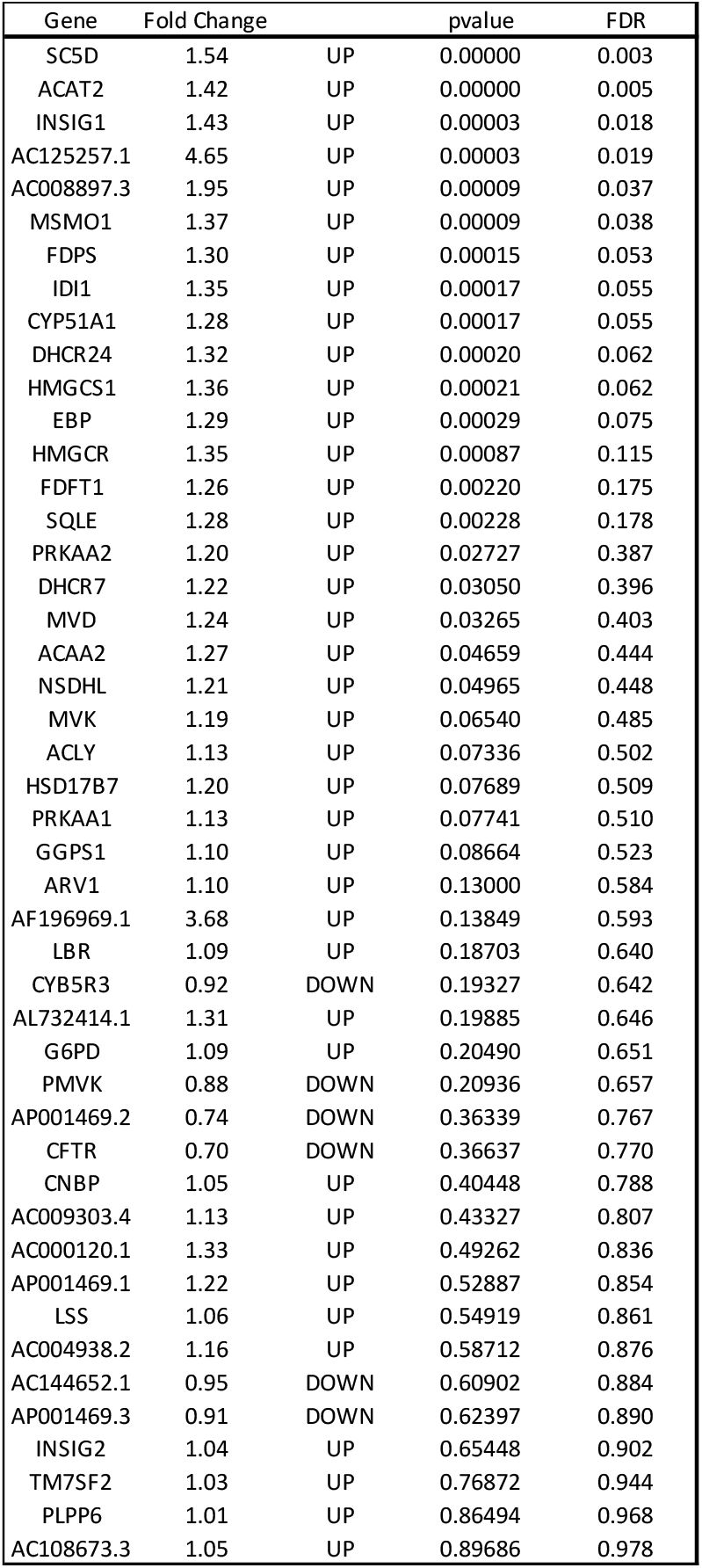
Results on all chlesterol genes

**Figure 2:**
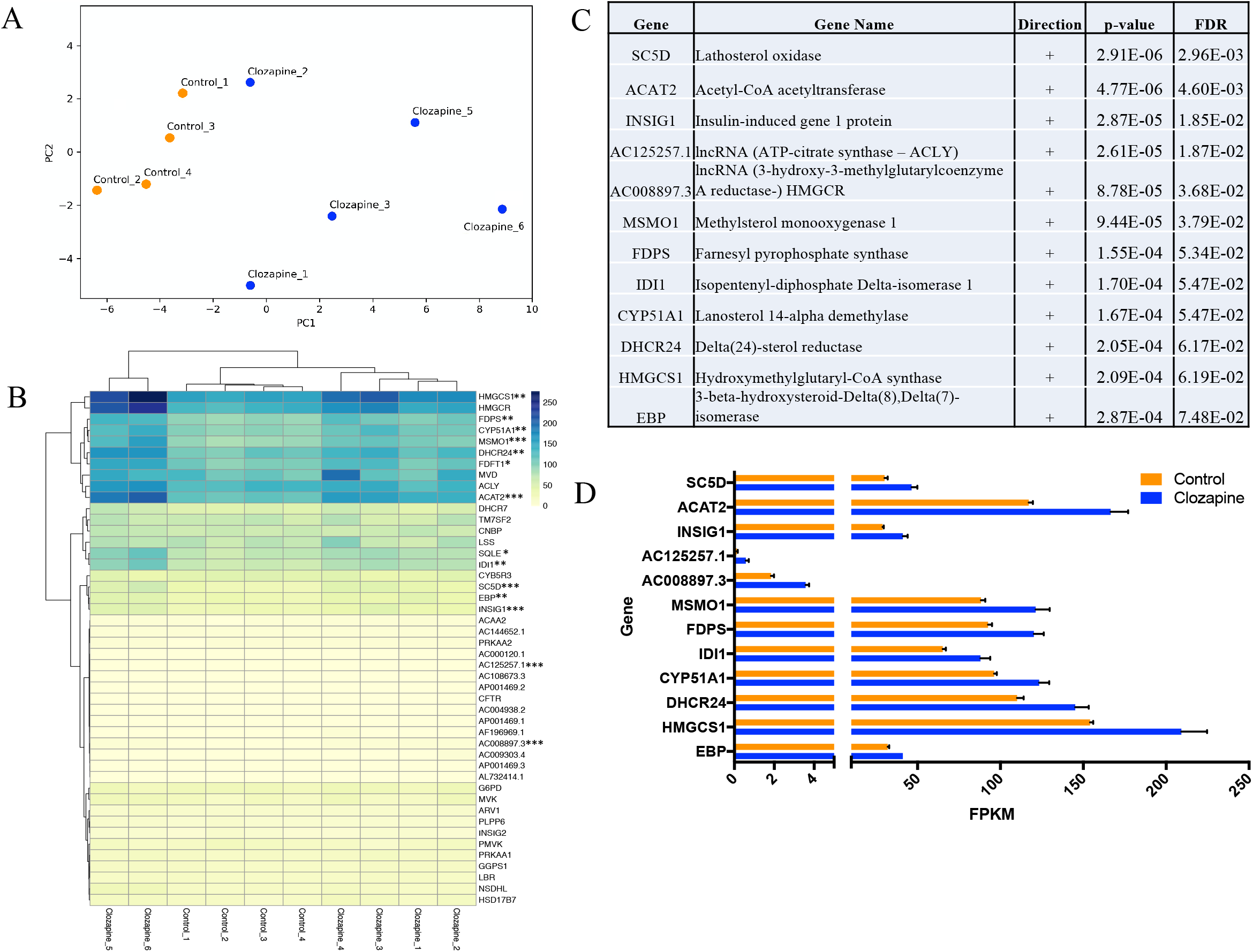
(A) Principal component Analysis of RNA-seq of clozapine-exposed IPSC-derived neurons. Control samples are represented in Orange. Clozapine-exposed samples are in Blue. (B) Dendrogram of Top 46 genes. Asterisks denotes significantly differentially expressed genes after Clozapine treatment according to FDR levels: *** for FDR 0.05, ** for FDR 0.1, * for FDR 0.2. Analysis with FDR cut-off of 0.05, 0.1, and 0.2 contains 6 genes, 12 genes, and 14 genes, respectively. (C) Top 12 significantly differentially expressed genes after Clozapine treatment. All top 12 genes are key components of Cholesterol Biosynthesis pathway with FDR<0.1. Expressivity level of top 12 genes. Bar graph shows expression level of each gene in pairs of treated (Blue) vs untreated IPSC-derived samples (orange). Error bars with **SEM**.

## Discussion

Multiple studies, most working with glial or glial-like cells, have found biosynthesis and regulation of fatty acids and cholesterol to be a common effect of antipsychotics (Bae and Paik, 1997; Ferno et al., 2005). In fact, it has been suggested that this could be one of the mechanisms through which these drugs ameliorate psychosis (Ferno et al., 2005; Polymeropoulos et al., 2009). The importance of cholesterol not only as a component of myelin but also as a key component of the lipid bilayer of the cellular membrane and a necessary component for synapse and dendrite formation (Goritz et al., 2005) is consistent with this hypothesis. Defects in brain cholesterol metabolism have been implicated in neurodegenerative diseases, such as Alzheimer’s disease, Huntington’s disease and Parkinson’s disease (Zhang and Liu, 2015). More importantly for psychiatric disease, Smith-Lemli-Opitz syndrome (SLOS) is caused by an inborn error of cholesterol biosynthesis and ~50% of individuals meet criteria for autism (Tierney et al., 2001).

Finally, the gene *HMCGR,* encoding the rate-controlling enzyme for the production of cholesterol and targeted by statins (Sirtori, 2014), was upregulated along with an associated lncRNA (AC008897.3). Interestingly, *HMCGR* has also been found to carry excess loss of function mutations in schizophrenia (meta-analysis p = 4.5×10^−4^, (schema.broadinstitute.org/results), directly implicating cholesterol metabolism as a potential contributor to the disease.

Induced pluripotent stem cells (iPSC) technology is a powerful tool for establishing *in vitro* models of disease. For neuropsychiatric disorders, this tool is particularly useful as live human neurons are essentially impossible to obtain. The feasibility of using iPSCs for disease modeling in SZ has been previously reported (Brennand et al., 2011; Chiang et al., 2011) and recently reviewed by Das et al. (Das et al., 2020). We used iPSC derived induced excitatory neurons to test their response to Clozapine, expecting it may be different than that of glial cells and highlight other possible mechanisms of antipsychotic function. To our surprise no pathway other than cholesterol metabolism emerged, while the robustness of the results confirmed that our isogenic in vitro model has substantial power. The Top 12 genes, all key components of the cholesterol biosynthesis pathway, include *SC5D, ACAT2, AC125257, INSIG1, AC008897.3, MSMO1, FDPS, CYP51A1, IDI1, DHCR24, HMGCR, HMGCS1, EBP*. Five of the top 12 genes have been previously reported to be upregulated by antipsychotics in human glioblastoma cells (*HMGCS1, FDPS, IDI1, CYP51* and *SCD*) (Ferno et al., 2005). Two of them and an additional three have been reported in human retinal pigment epithelia cell line ARPE-19 (*ACAT2, INSIG1, EBP* along with *FDPS, CYP51*) (Polymeropoulos et al., 2009).

In addition to the effects we observe on cholesterol metabolism Clozapine has other known functions on neurons including on dopamine and serotonin receptors. In drug-resistant schizophrenia where Clozapine is most often prescribed, abnormalities in the glutamatergic system may be particularly relevant (Aringhieri et al., 2018) and therefore our study of glutamatergic neurons seems appropriate. However, studying one cell type in isolation cannot capture the complexity of interactions between neuronal populations in the brain and in this case will miss the effects on dopamine or serotonin receptors, which is currently unavoidable but still a significant limitation of our study.

It is remarkable though in view of the very strong results on cholesterol that no other functional category reached statistical significance. By singling out cholesterol metabolism as the main response to antipsychotics in glutamatergic neurons, as previously reported on glial and retinal cells, our work provides support for the hypothesis that this may be a major mechanism in the therapeutic benefits of Clozapine. Understanding all possible mechanism that can contribute to disease improvement is an important component to future tailored treatments guided by precision medicine.

## Conclusions

There have been several studies suggesting that antipsychotics have bilayer-modifying potencies. Our experiment singles out biosynthesis and regulation of fatty acids and cholesterol metabolism as the main response to Clozapine in human neurons consistent with the previous reports on glial cells. This finding suggests that alterations in lipid homeostasis in dopaminergic neurons might underlie the pathogenesis of schizophrenia and adds support for the hypothesis that this may be a major mechanism in the therapeutic benefits of Clozapine and possibly other antipsychotics.

## Acknowledgement

This work was supported in part by NIMH grants P50 MH094268, R01 MH113215 and RF1 MH122936 to DA.

**Bio-samples and/or data for this publication were obtained from NIMH Repository & Genomics Resource, a centralized national biorepository for genetic studies of psychiatric disorders.**

Author contributions: D.D. performed all laboratory experiments. X.P. performed data analysis guided by J.B. and D.A. A.L. and D.A. prepared the manuscript with input from all other authors. D.A and D.D. designed the project.

This study was approved by the Johns Institutional review board (protocol IRB00122135).

